# Endogenization from diverse viral ancestors is common and widespread in parasitoid wasps

**DOI:** 10.1101/2020.06.17.148684

**Authors:** Gaelen R. Burke, Heather M. Hines, Barbara J. Sharanowski

## Abstract

The Ichneumonoidea (Ichneumonidae and Braconidae) is an incredibly diverse superfamily of parasitoid wasps that includes species that produce virus-like entities in their reproductive tracts to promote successful parasitism of host insects. Research on these entities has traditionally focused upon two viral genera *Bracovirus* (in Braconidae) and *Ichnovirus* (in Ichneumonidae). These viruses are produced using genes known collectively as endogenous viral elements (EVEs) that represent historical, now heritable viral integration events in wasp genomes. Here, new genome sequence assemblies for eleven species and six publicly available genomes from the Ichneumonoidea were screened with the goal of identifying novel EVEs and characterizing the breadth of species in lineages with known EVEs. Exhaustive similarity searches combined with the identification of ancient core genes revealed sequences from both known and novel EVEs. Two species harbored novel, independently derived EVEs related to a divergent large double-stranded DNA (dsDNA) virus that manipulates behavior in other hymenopteran species. While bracovirus or ichnovirus EVEs were identified as expected in three species, the absence of ichnoviruses in several species suggests that they are independently derived and present in two younger, less widespread lineages than previously thought. Overall, this study presents a novel bioinformatic approach for EVE discovery in genomes and shows that three divergent virus families (nudiviruses, the ancestors of ichnoviruses, and LbFV-like viruses) are recurrently acquired as EVEs in parasitoid wasps. Virus acquisition in the parasitoid wasps is a common process that has occurred in many more than two lineages from a diverse range of arthropod-infecting dsDNA viruses.

**Significance:** Parasitoid wasps are an extremely diverse group of animals that are known to harbor Endogenous Virus Elements (EVEs) that produce virions or virus-like particles of key importance in wasps’ parasitism success. However, the prevalence and diversity of independently acquired EVEs in parasitoid wasp lineages has remained largely uncharacterized on a widespread scale. This study represents an important first step and hints at the massive, untapped diversity of EVEs in parasitoid wasps via the identification of several novel virus co-option events from diverse groups of double-stranded DNA virus pathogens.

## Introduction

Although viruses have long been viewed as pathogenic organisms, there is now ample evidence that viruses can confer important benefits to their hosts and have played a major role in the evolution of life on earth (Goic & Saleh 2012; Villarreal & Witzany 2010; Schmitt & Breinig 2002; Ryabov et al. 2009; Rossignol et al. 1985; Barton et al. 2007; Brown et al. 2006; Malmstrom et al. 2005; Moran et al. 2005; Strand & Burke 2020; Dunlap et al. 2006). Further, in most sequenced eukaryotic genomes there is evidence of viral gene “footprints” (Feschotte & Gilbert 2012), suggesting viral endogenization is common and could play an important role in genome and organismal evolution. Understanding how beneficial viral associations evolve is essential for a holistic view on the evolution of eukaryotic organisms (Roossinck 2011; Villarreal 2015).

Perhaps some of the most complex examples of viral endogenization occur in parasitic wasps belonging to the exceptionally diverse superfamily Ichneumonoidea (Braconidae + Ichneumonidae) with more than 44,000 described species (Yu et al. 2012). As parasitoids, these wasps lay their eggs in or on other insect “hosts”, where their progeny feed and complete the immature stages of development, resulting in the death of the host. Conservative estimates suggest that one in ten animals is a parasitoid wasp (Askew 1971) and more recent estimates exceed these already astonishing numbers (Forbes et al. 2018; Jones et al. 2009; Rodriguez et al. 2013). Among parasitoid wasps, Ichneumonoidea are particularly diverse, comprising ~28% of all Hymenoptera, ~3% of all terrestrial, multicellular life (Chapman 2009; Yu et al. 2012) and parasitizing a broad range of insects and other arthropods. To facilitate host invasion, ichneumonoid wasps are known to employ novel viral associations in the form of Endogenous Viral Elements (EVEs), in which elements of viral genomes become permanently integrated into the genomes of wasps (Bezier et al. 2009; Burke, Simmonds, et al. 2018; Pichon et al. 2015; Volkoff et al. 2010; Béliveau et al. 2015). EVEs produce virions or virus-like particles (VLPs) in ovaries of these wasps, which are injected into hosts during parasitism. Based upon experimental studies in a number of representative species, it is thought that these viruses function in the promotion of successful parasitism (Edson et al. 1981; Beckage et al. 1994; Rotheram & Salt 1973; Salt 1965) and may contribute to the immense diversity of these parasitic wasps. Viruses or VLPs function in the delivery of virulence molecules such as DNA or proteins that suppress host immune defenses or alter host development and behavior in ways that facilitate survival, growth and development of the larval parasitoid (Strand 2012; Darboux et al. 2019; Lee et al. 2009; Reineke et al. 2006). Remarkably, many if not all viruses or VLPs produced by wasps stem from pathogenic viral ancestors (Pichon et al. 2015; Bezier et al. 2009; Burke, Simmonds, et al. 2018; Stasiak et al. 2005).

Polydnaviruses (PDVs, belonging to the family Polydnaviridae, Strand & Drezen 2012) are EVEs documented to occur within select but diverse clades within Ichneumonoidea. In Braconidae, they are present in the microgastroid complex (*sensu* Sharanowski et al. 2011) comprising at least 6 subfamilies with more than 3900 described species. In Ichneumonidae, PDVs have been discovered in two families: Campopleginae and Banchinae, with more than 2,200 and 1,750 described species, respectively (Yu et al. 2012). Most large double-stranded DNA viruses have genes that can be divided into two categories: 1) replication genes, which encode essential replication machinery and are conserved among genomes, and 2) virulence genes of diverse origins whose products interact with host defenses and are gained and lost much more rapidly (Yutin et al. 2009; Kawato et al. 2018; Rohrmann 2011). PDVs have two components that are dispersed within the genomes of wasps: replication genes and proviral segments (the regions of the genome that are packaged into virions and contain virulence genes). The replication machinery for these viruses is not packaged into virions, making PDVs replication-defective and thus reliant on the wasp for replication (Strand & Burke 2014; Burke et al. 2014; Bézier et al. 2009; Bézier et al. 2013; Volkoff et al. 2010). The inheritance of permanently integrated PDVs is predicted to produce genetically unique, but related EVEs in each wasp species within the three major clades of PDV-carrying wasps (Whitfield & Asgari 2003).

Despite such strong functional similarity, evidence shows that known PDVs have at least two unique origins. The family Polydnaviridae is divided into two genera: *Bracovirus* and *Ichnovirus* (Strand & Drezen 2012). The morphology of the polydnaviruses found in Ichneumonidae (ichnoviruses) and Braconidae (bracoviruses) are vastly different (Stoltz & Whitfield 1992), and genomic sequences of the replication machinery show affinity to different classes of viruses (insect beta-nudiviruses and related baculoviruses for bracoviruses; relatives of nucleocytoplasmic large DNA viruses (NCLDVs) for ichnoviruses) (Béliveau et al. 2015; Bezier et al. 2009; Volkoff et al. 2010). In the Braconidae, bracoviruses have been localized histologically in species across the monophyletic microgastroid complex (Whitfield, 1997), suggesting a single PDV origin in this clade ~100 Mya (Murphy et al. 2008).

In addition to these PDVs, several types of EVEs have been recently discovered within Ichneumonoidea. For example, in the ichneumonid *Venturia canescens* (Campopleginae), viral replication genes were co-opted from the insect alpha-nudiviruses (related but distinct from the PDV progenitors, the beta-nudiviruses) and used to produce VLPs in wasp ovaries (Pichon et al. 2015). The *V. canescens* genome lacks the genes required to make a capsid to house viral DNA in virions, preventing delivery of virulence genes but allowing delivery of wasp-derived virulence proteins into hosts within VLPs (Feddersen et al. 1986; Pichon et al. 2015). Through wasp genome sequencing, an independent acquisition of viral genes from the alpha-nudiviruses was recently identified in *Fopius arisanus* (Braconidae), the first recognized incidence of viral genome integration in the subfamily Opiinae (Burke, Simmonds, et al. 2018). Other reports of reproductive gland associated viruses in the Ichneumonoidea have been published (>35), but are limited to the identification of virions in wasp tissues and do not include any genetic analyses (Suzuki & Tanaka 2006; Lawrence 2005). Two very recent studies have documented the presence of EVEs outside of the Ichneumonoidea in parasitoid species belonging to the Figitidae and Chalcididae (Di Giovanni et al. 2020; Zhang et al. 2020). Recently discovered EVEs have not been assigned to Polydnaviridae because thus far no VLPs produced package DNAs and because the family is polyphyletic and likely to be revised in the future. These data suggest that viral co-option events may be more common in parasitoids than previously thought.

Thus far, genetic discovery of these EVEs has been sporadic. Comprehensive exploration of the number of integration events and the rules governing their acquisition and function requires improvements in genomic pipelines for viral identification. In this study, we seek to better understand the diversity of viral origins in ichneumonoid wasps through developing a comparative genomic approach for EVE discovery. Using new genome sequence datasets in combination with publicly available genome assemblies from species belonging to Ichneumonoidea, this research has three objectives: 1) to develop a method for the identification of endogenous virus elements derived from diverse large double-stranded DNA viruses; 2) to examine the breadth of PDV incidence in lineages known to produce bracoviruses and ichnoviruses; and 3) to identify novel EVEs likely to produce virions or VLPs in species in which they have not been described previously. Screening of 17 genomes from parasitoids within the Ichneumonoidea (including eleven new genome assemblies) identified both familiar and novel EVEs in a number of species. These results challenge existing assumptions about the species distribution and origins of ichnoviruses, and find a new family of double-stranded DNA viruses that are common EVE progenitors.

## Materials and methods

### Wasp species sampling

Sharanowski et al. (2020) recently generated genome sequence datasets from 11 species for the purpose of resolving the phylogenetic relationships among species in the Ichneumonoidea. The sampling included seven species from Ichneumonidae and four species from Braconidae, all derived from different subfamilies to maximally capture diversity (Table 1, Supplementary Table 1). All DNA samples used for sequencing were derived from single adult females, except *Odontocolon* sp, which was sequenced from a single adult male. Additionally, six publicly-available genomes from the Ichneumonoidea (five braconid and one ichneumonid species) with known presence or absence of viral associations served as controls for this study (Table 1, Burke, Walden, et al. 2018; Shi et al. 2019; Tvedte et al. 2019; Geib et al. 2017; Leobold et al. 2018; Yin et al. 2018).

**Table 1.** Taxa used for genome analysis. Higher Group placement is taken from Sharanowski et al. (2011) and Sharanowski et al. (2020). Taxa not identified to species due to difficulty in accurate identification are listed as sp.

| Assembly accession | Family | Higher Group | Subfamily | Genus species | Expected or known viral association |
| --- | --- | --- | --- | --- | --- |
|  | Braconidae | Microgastroid | Cheloninae | *Phanerotoma* sp. | Bracovirus |
|  | Braconidae | Euphoroid | Euphorinae | *Meterous* sp. | Unknown |
|  | Braconidae | Helconoid | Helconinae | *Eumacrocentrus americanus* | Unknown |
|  | Braconidae | Cyclostomes s.s. | Rogadinae | *Aleiodes* sp. | Unknown |
|  | Ichneumonidae | Ichneumoniformes | Adelognathinae | *Adelognathus* sp. | Unknown |
|  | Ichneumonidae | Ophioniformes | Banchinae | *Lissonota* sp. | Ichnovirus |
|  | Ichneumonidae | Ophioniformes | Campopleginae | *Dusona* sp. | Ichnovirus |
|  | Ichneumonidae | Ophioniformes | Ctenopelmatinae | *Anoncus* sp. | Unknown |
|  | Ichneumonidae | Ophioniformes | Mesochorinae | *Mesochorus* sp. | Unknown |
|  | Ichneumonidae | Pimpliformes | Pimplinae | *Dolichomitus* sp. | Unknown |
|  | Ichneumonidae | Xoridiformes | Xoridinae | *Odontocolon* sp. | Unknown |
| Previously published accessions |  |  |  |  |  |
| GCA_000956155.1 | Braconidae | Microgastroid | Microgastrinae | *Cotesia vestalis* | Bracovirus |
| GCA_001412515.3 | Braconidae | Alysoid | Opiinae | *Diachasma alloeum* | None discovered |
| GCA_000806365.1 | Braconidae | Alysoid | Opiinae | *Fopius arisanus* | Endogenous nudivirus |
| GCA_002156465.1 | Braconidae | Macrocentroid | Macrocentrinae | *Macrocentrus cingulum* | None discovered |
| GCA_000572035.2 | Braconidae | Microgastroid | Microgastrinae | *Microplitis demolitor* | Bracovirus |
| N/A* | Ichneumonidae | Ophioniformes | Campopleginae | *Venturia canescens* | Endogenous nudivirus |

\* Genome available at: http://bipaa.genouest.org/sp/venturia_canescens

### Genome assembly

Illumina sequence reads generated from genomic DNAs from 11 wasp species (PE100) were screened with trimmomatic v.0.36, using the parameters “LEADING:3 TRAILING:3 SLIDINGWINDOW:4:15 MINLEN:36”. Any read pairs with overlapping 3’ ends were merged with pear v.0.9.8 using default settings. Paired reads, merged reads, and unpaired single reads were used as input for *de novo* assembly of contigs using SPAdes v.3.12.0 (Bankevich et al. 2012) with kmers of 21, 33, 55, 77 bp and default parameters.

### Database construction and rapid homology searches for curation of assemblies and identification of EVEs

A custom database was made to identify contigs that contain genes of viral origin and exclude contaminant contigs (Medd et al. 2018). The database contained all protein sequences from the NCBI refseq database (downloaded February 2019) from Hymenoptera, Lepidoptera, Diptera, Coleoptera, bacteria, archaea, nematodes, and fungi, and viral protein sequences from the NCBI nr database (downloaded February 2019). TaxonKit v.0.3.0 was used to generate lists of all NCBI taxonomy ID numbers for included groups of organisms (Shen & Xiong 2019). The software csvtk v.0.15.0 (https://bioinf.shenwei.me/csvtk/) was used to obtain accession numbers from proteins derived from species with included taxonomy IDs using the NCBI database prot.accession2taxid.gz downloaded in February of 2019. Protein sequences from included species were then extracted from the refseq and nr databases and made into a database with diamond v.1.0 (Buchfink et al. 2015). Sequences from hymenopteran genomes with known EVEs were excluded so that any genes with viral origin would have hits to viral proteins and be assigned to the correct taxonomic lineage (virus rather than insect). Genes from complete polydnavirus genomes (proviral genome segments) were also excluded from the database because they do not contain viral replication genes and do contain many genes of eukaryotic origin. The final database contained 117,461,186 protein sequences.

ORFs were identified from genomic contigs (generated as described below) or scaffolds using emboss v.6.6.0 getorf with minimum size of 150bp yielding between 247,340 and 691,147 ORFs per genome assembly (Rice et al. 2000). ORFs were searched against the database described above with diamond with an e-value threshold of 0.01, retaining a single top hit. NCBI taxonomy IDs were included in diamond output reports.

### Removal of sequence contaminants and assessment of assemblies

Contigs smaller than 200bp in size were removed from final assemblies. Blobtools v1.1 (Laetsch & Blaxter 2017), was used to retain only those contigs that were assigned to Arthropoda and Viruses or that had no hits. Assembly statistics were evaluated using quast v.5.0.2 (Gurevich et al. 2013). The completeness of each set of genome sequence contigs was analyzed by identifying the number of arthropod Benchmark Universal Single-Copy Orthologs (BUSCOs) (Simão et al. 2015). BUSCO v.3.0.2 was run on the assembled contigs or genome scaffolds for previously published genomes (“-m geno”) to identify orthologs in the Insecta ortholog database version 9, using *Nasonia* models for gene prediction.

To determine the read coverage of assembled contigs, all reads used for sequence assembly were mapped to the contigs using bowtie2 v.2.3.4.1, and coverage calculated with samtools bedcov v.1.9 (Langmead & Salzberg 2013; Li et al. 2009). A similar process was used to determine read coverage of genome scaffolds in previously published genomes, except that sequence reads were downloaded from the NCBI Sequence Read Archive (SRA) when available and processed using trimmomatic and pear as described above.

### Identification of genes of viral origin in wasp genomes

Identification of virus-derived genes in wasp genomes was done using several approaches. All eleven new and previously published genome assemblies were searched without any *a priori* expectation for the presence of virus-derived genes by: 1) exhaustively screening all ORFs from genome assemblies for viral hits against a curated database; and 2) searching all ORFs for matches to ancient core genes found in large double-stranded DNA virus groups that are known to infect insects (see below). These methods were expected to non-exhaustively discover the presence of virus-derived genes to narrow down the list of species for which more detailed identification and annotation should be performed.

### i. Exhaustive screening of ORFs for viral hits

The first approach used was modified from Medd et al. 2018, and used exhaustive similarity searches of all possible Open Reading Frames (ORFs) (of a minimum size) against a curated database to identify sequences with similarity to viral proteins. Despite the fragmented state of the newly added genome assemblies, it was reasoned that the majority of genes of viral origin and architecture would be intact (not be broken into pieces across contigs) because N50 values were equal to or greater than the expected sizes of these genes (less than 1000 base pairs on average). Lineage information was generated by TaxonKit and allowed for categorization of hits by virus type and family.

### ii. Targeted searches for ancient core genes in dsDNA viruses

In the second approach, the set of ORFs with viral hits from each wasp species were then queried for the presence of ancient core genes. In order to do this, Hidden Markov Models were constructed for six ancient core genes (*DNApol, helicase, lef-5, lef-8, lef-9*, and *p33*, Wang et al. 2012) to find deeply divergent matches. Protein sequences were collated for 24 representative viral species: AcMNPV, CpGV, NeseNPV, CuniNPV, ToNV, OrNV, HzNV-2, PmNV, DiNV, GbNV, GpSGHV, MdSGHV, LbFV, WSSV, CoBV, AMEV, MSEV, VACV, IIV-6, LDV1, HvAV-3e, PbCV1, ApMV, and HHV-3. Sequences were aligned using MUSCLE v3.8.31 (for use as PSSMs in PSI-BLAST) and made into HMMs using hmmbuild within HMMER v3.1b1 (Eddy 2011; Edgar 2004). HMMs were searched against ORFs with viral hits from each species using hmmsearch. The dispersed nature of genes in all parasitoid EVEs discovered to date make it likely that at least some virus-derived genes would be recovered, even though some genomes were poorly assembled (Supplementary Table 3).

### iii. Detailed annotation of EVEs using homology to other EVE sequences

If sets of virus-derived genes were confidently identified in a wasp genome assembly using the above methods, more detailed annotation was employed for these species. *Post hoc* searches of wasp genome assembly ORFs against the previously excluded replication genes from bracovirus, ichnovirus, and other parasitoid EVEs were employed to further identify virus-derived genes that may not have been found in earlier searches due to sequence divergence from viral ancestors. To effectively identify virus-derived genes in wasp genomes expected to contain bracoviruses or ichnoviruses, several publicly available reference datasets were employed. The genes encoding the nudivirus-like structural components of bracovirus virions have been completely catalogued in *Microplitis demolitor* and partially in *Cotesia congregata* (both subfamily Microgastrinae ~53myo) and *Chelonus inanitus* (subfamily Cheloninae, ~85myo) (Murphy et al. 2008; Bezier et al. 2009; Burke et al. 2014). The genes contributing to the production of ichnoviruses known as Ichnovirus Structural Protein Encoding Regions (IVSPERs) have been identified previously in wasps from the subfamily Campopleginae (*Hyposoter didymator*) and the subfamily Banchinae (*Glypta fumiferanae*) (Béliveau et al. 2015; Volkoff et al. 2010). Protein sequences from nudivirus-like replication genes from *M. demolitor, C. congregata*, and *C. inanitus*, and separately, protein sequences from IVSPERs in the ichnovirus-producing wasp species *Hyposoter didymator* and *Glypta fumiferanae* were added to the custom diamond database described above. All ORFs from each species were searched against the modified database (as above), and ORFs with top hits to sequences from bracovirus-producing or ichnovirus-producing wasp species were extracted from the output. Finally, when genes related to similar virus pathogens were found in multiple wasp genomes, these genes and surrounding ORFs were searched among the multiple wasp genomes to further identify and annotate virus-related genes.

### iv. Metrics used to determine whether virus-derived sequences are endogenous

Virus-derived sequences identified in wasp genome assemblies could be indicative of EVEs or “contamination” by sequences derived from an actively replicating viral infection. The latter would have a self-contained viral genome, ie. contigs could be assembled into a single, circular or linear dsDNA chromosome representing the viral genome of a non-integrated infectious virus. Features of genome architecture and sequencing metrics were used to build support for the hypothesis that virus-derived sequences are integrated into a given wasp genome. First, it was determined whether virus-derived genes were interspersed among non-viral genes in the wasp genome. Virus-derived genes with “viral architecture” were defined as genes that had homology to genes in viral genomes, are short and most often intronless. Non-viral genes were defined as having homology to hymenopteran or other animal genes and had “eukaryotic architecture”; comprising longer genes most often containing introns. If novel virus-derived genes were identified in a wasp genome assembly, scaffolds upon which viral genes were identified were annotated with MAKER v. 3.01.02-beta for eukaryotic genes (Holt & Yandell 2011) and Prokka v. 1.12 for virus-derived genes (Seemann 2014) to see if there were clear signatures of flanking genes with eukaryotic architecture. If available, sequence expression data were aligned to gene annotations with GMAP v.2019-05-12 (Wu & Watanabe 2005) to determine whether they were expressed and represent high-confidence annotations. Second, unless extremely recently derived, EVEs are expected to be present in all individuals and populations of a given species of wasp. If available, different versions of wasp genome assemblies generated from geographically distinct populations of wasps were queried for homologous viral genes with BLAST (Altschul et al. 1997). Third, characteristics of contiguous blocks of virus-derived genes were used to assess whether they were likely to be derived from EVEs or an actively-replicating virus infection. Most core replication genes are present as a single copy in insect viral pathogens with dsDNA genomes (Yutin et al. 2009) whereas degraded, pseudogenized forms of viral genes can be found in genomes with EVEs. Finally, sequence read coverage and GC content of contigs were used as further lines of evidence for differentiation of endogenous vs. replicating viral sequences. As EVEs are integrated into wasp genomes, the sequence read coverage of contigs containing EVEs is expected to be similar to other contigs. However, high coverage could be observed from local chromosomal amplification of EVEs, which is known to occur in several parasitoid wasps (Burke et al. 2015; Louis et al. 2013; Di Giovanni et al. 2020). Coverage that is higher or lower than wasp genome contigs could also be an indication of the presence of viral DNA contaminating the insect DNA sequenced. EVEs that represent ancient integration events (such as bracoviruses, Burke & Strand 2012) are expected to have a GC content similar to the remainder of the wasp genome, while differing GC content between contigs with virus-derived genes and others could be indicative of recently acquired EVEs (whose ancestors may have had differing GC content) or actively replicating viral contaminants. These metrics were used when available and in conjunction with each other to provide an overall picture of the likelihood that virus-derived genes represent EVEs.

### Phylogenetic analyses to determine the closest known relatives of EVEs

To make single gene trees, amino acid sequences from ORFs with viral hits and from endogenous viruses in wasp genomes were added to sequences from representative taxa used to make PSSMs and HMMs as above. BLAST against the NCBI nr database was used to identify any additional endogenous viral genes from parasitoid wasps to include in alignments. Protein sequences from the EVEs in *Glypta fumiferanae* and *p33* sequences from nimaviruses were excluded because they were extremely divergent and decreased alignment quality. Sequences were aligned using the MAFFT einsi algorithm (Katoh & Standley 2013), and maximum likelihood phylogenies were derived using the RAxML with the PROTGAMMALG model using default parameters and 100 bootstrap replicates (Stamatakis 2014).

A multi-gene phylogeny of NALDV representatives was generated by aligning protein sequences with MUSCLE (Edgar 2004), concatenated with FASconCAT-G (Kück & Longo 2014) and trimmed with default parameters and a gap threshold of 0.6 by TrimAl (Capella-Gutierrez et al. 2009). A maximum likelihood phylogeny was constructed for the concatenated alignment using RAxML with default parameters and 1,000 rapid bootstrap replicates in the CIPRES portal (Miller et al. 2010).

## Results

### Sequencing and assembly generated 11 new draft genome sequences for parasitoid wasps

To accomplish the objective of identifying known and novel EVEs in parasitoid genomes, it was necessary to screen new genomic data for a range of wasp species that had both expected and unknown presence of EVEs (Table 1, Figure 1). In the literature, it is taken for granted that EVEs are present within entire clades of parasitoid species (for example, the microgastroid lineage of Braconidae or the Campopleginae or Banchinae subfamilies of the Ichneumonidae), which sets up expectations for the identification of EVEs in particular species with newly-generated genome assemblies (Table 1, Figure 1). For newly sequenced taxa, it was expected that bracovirus genes would be found within *Phanerotoma sp*. (subfamily Cheloninae in the microgastroid lineage) and ichnovirus genes (e.g. IVSPER genes) within *Lissonota sp*. (Banchinae) and *Dusona sp. 2* (Campopleginae, hereafter *Dusona sp*). It was unknown whether the remaining eight species would have EVEs in their genomes.

**Figure 1.**
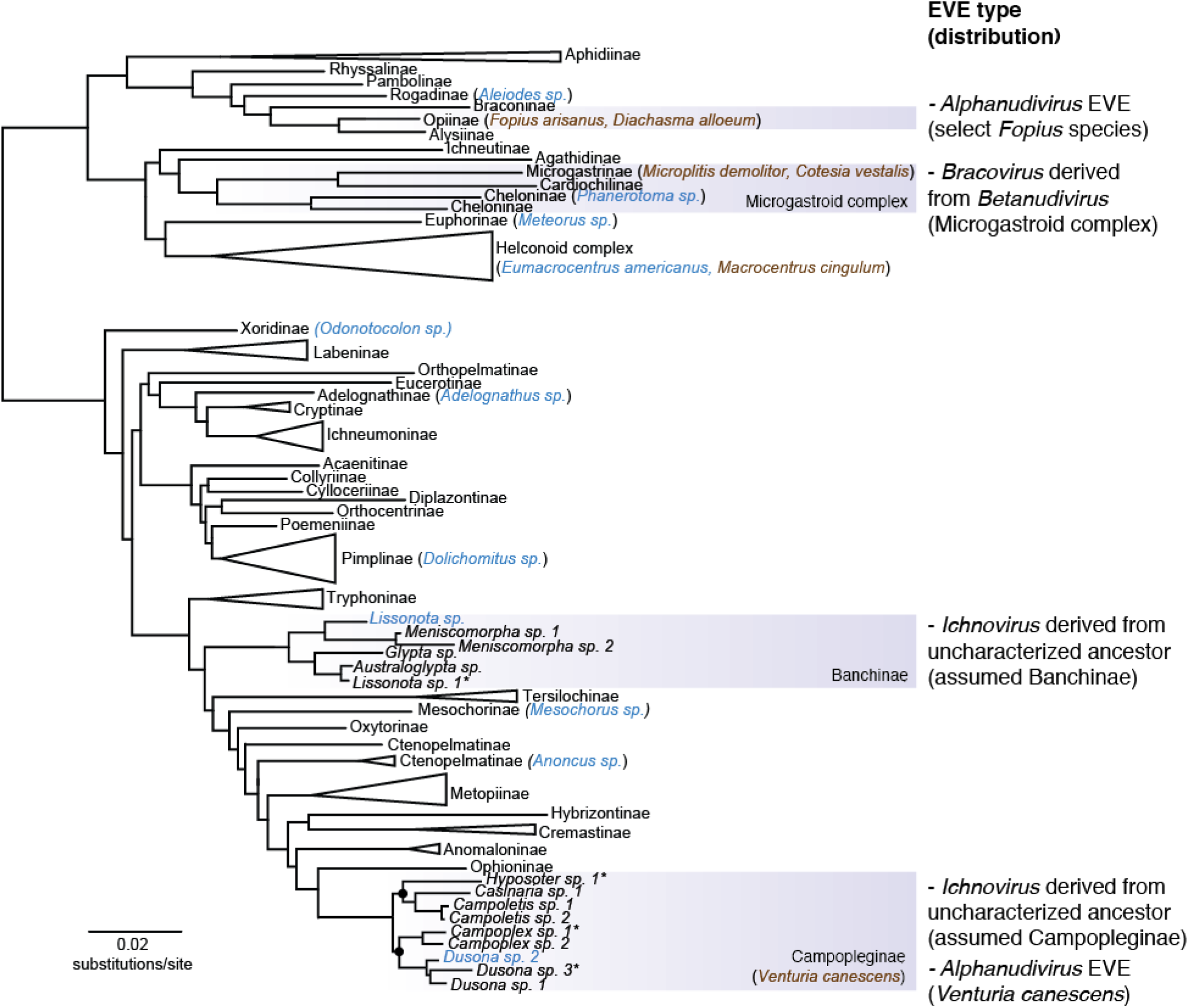
Maximum likelihood tree of Ichneumonoidea from Sharanowski et al. (2020). Subfamilies with multiple representatives (except the microgastroid complex, Campopleginae, and Banchinae) have been collapsed for viewing subfamily relationships, the presence and distribution of EVEs, and representative species analyzed in this study. Subfamilies with previously published, genetically characterized EVEs are shaded in boxes, and to the right, independent origins of EVEs, the type of virus ancestor they are derived from, and their known or assumed distribution is indicated. Two nodes within the Campopleginae are highlighted showing that the subfamily is divided into two major clades. Species analyzed in this study are placed next to the subfamily to which they belong and are colored blue (new genome assemblies) and brown (previously published genome assemblies). Taxa marked with an asterisk after the name have uncertain species identification. Support for the tree is robust and can be viewed in detail in Sharanowski et al. (2020).

Illumina read counts from the 11 newly added genomes ranged from 58.9 – 88.7 million per library (Supplementary Table 2). Given that the sequence data were generated from short insert libraries only, the assemblies yielded many contigs (ranging from 13,843 to 423,888) with relatively low N50 values (1,110 to 31,843 nucleotides) compared to the current standard for complete insect genome sequences that employ scaffolding with large insert sequencing libraries or long-sequence read technologies (Supplementary Table 1). The cumulative sizes of scaffolds (a proxy for genome size) was similar for new and previously published high quality assemblies. BUSCO analysis revealed variation in the level of completeness of each new genome assembly, ranging from detection of 98.7% complete BUSCOs in *Meteorus* to detection of 26.8% complete and 46.4% fragmented BUSCOs (total 73.2%) in *Anoncus* (Supplementary Table 3). Although some genome assemblies were relatively fragmented, overall the amount of sequence data was likely enough to cover a majority of the genomes of these wasps.

### Identification of genes of viral origin using exhaustive sequence similarity searches is fraught with false positives

To identify genes responsible for making virions or VLPs in wasp genomes, a method to identify endogenous virus elements (EVEs) derived from diverse large double-stranded DNA viruses was needed. Although actively replicating viruses and lateral transfers of single genes from viruses into eukaryotic hosts can generate novel phenotypes of key importance (Parker & Brisson 2019; Coffman et al. 2020; Aswad & Katzourakis 2012; Dunlap et al. 2006; Lavialle et al. 2013), the focus here is upon identification of sets of viral replication genes that could be responsible for the production of virions or VLPs that are permanent key components of wasp biology and parasitism success. Homology searches of all ORFs extracted from genome assembly scaffolds against a custom database produced viral hits for 37 to 75 ORFs for species with previously published genomes, and 10 to 53 ORFs for new species (Supplementary Figure 1 and 2, Supplementary Table 4). Hits were to viruses belonging to several major groups: dsDNA viruses, ssDNA viruses, ssRNA viruses, dsRNA viruses, Ortervirales, and unclassified viruses. As all genetically characterized examples of virions or VLPs produced by parasitoid wasps thus far originate from large dsDNA viruses, hits to other types of viruses were not analyzed further in this study. Most hits were to the dsDNA viruses, particularly to the families Nudiviridae, Baculoviridae, Poxviridae, Ascoviridae, Hytrosaviridae, Iridoviridae, and the Caudovirales. However, manual inspection of the annotations associated with these hits revealed that many of these proteins are retroviral proteins, inhibitor of apoptosis proteins, chitosanases, and transposases (Supplementary Table 4). These genes do not encode the necessary components for building virions or virus-like particles, and are frequently transferred between viruses and eukaryotes. Therefore, many or most of these genes have obscure origins and are unlikely to play a role in producing virions or VLPs that contribute to wasp parasitism success.

### Targeted searches for ancient core genes can accurately identify EVEs likely to produce virions or VLPs in wasp genomes

The preceding results indicated that a different approach was necessary to identify sets of virus-derived genes that are likely to be involved in producing virions or virus-like particles. Double-stranded DNA viruses that infect insects can be categorized into several virus families belonging to at least two major groups, the monophyletic Nuclear Cytoplasmic Large DNA Viruses (NCLDVs) and Nuclear Arthropod-specific Large DNA Viruses (NALDVs) (Iyer et al. 2001; Wang et al. 2012). While viruses generally use a very diverse set of strategies for replication, there exist at least six genes encoding replication components that are common to NALDVs and NCLDVs (Wang et al. 2012). These genes are present in most extant members of major families of viruses in these groups and are referred to as ancient core genes (although they may not share a common origin, Iyer et al. 2006; Wang et al. 2012).

To produce the key components necessary for the completion of virus replication and construction of virions or VLPs, it was rationalized that wasp genomes would need to contain at least some of the ancient core genes. To identify the presence or absence of ancient core genes, each set of ORFs with viral hits generated by the exhaustive method above was searched with methods that can detect homologs with very low sequence similarity by focusing upon patterns of conserved sites among protein sequences. While genes encoding ichnovirus replication machinery may be related to NCLDV core genes, the protein sequences from IVSPERs were too divergent to be incorporated into our search strategies. Thus, searches for ancient core genes are expected to identify EVEs derived from NCLDVs and NALDVs, but not the ancestors of ichnoviruses.

PSI-BLAST and HMM searches among putative viral protein sequences from each wasp species identified a total of 11 homologs of ancient core genes in new genome assemblies and 22 in previously published genome assemblies (Table 2, Supplementary Table 5). Previously published genome sequences known to contain EVEs served as positive controls for the identification of ancient core genes using the method described here that relies upon limited homology from deep divergence events. Previous work involving manual annotation of EVEs in the *M. demolitor, V. canescens*, and *F. arisanus* genomes indicated that all contain five out of six of the ancient core genes (a viral DNA polymerase gene is often lost in wasp species with EVEs, Burke 2019). Homology searches identified genes encoding *helicase, lef-8*, *lef-9* and *p33* proteins in the set of putative viral proteins for each positive control species. However, *lef-5* was identified in *F. arisanus* only, and could not be identified in the two other species even when the full set of ORFs from each species was searched. This may be due to limited sequence similarity and the short sequence length of *lef-5* genes reducing the success of PSI-BLAST and HMMR searches targeting very diverse taxa. Although *C. vestalis* females are known to produce bracovirus in their ovaries, no publication has yet provided a detailed annotation of EVEs in the *C. vestalis* genome. ORFs with homology to viral *helicase*, *lef-8*, and *lef-9* genes were identified, as well as a viral DNA polymerase (*DNApol*), warranting further exploration of genes of viral origin in this species (see below). Two previously published wasp genomes that are not known to contain EVEs (*D. alloeum* and *M. cingulum*) were negative for all ancient core genes. These results indicate that PSI-BLAST and HMMR can reliably detect the presence or absence of viral ancient core genes in wasp genomes.

**Table 2.** Matches to ancient core genes identified in sequenced wasp genomes.

| Species | DNA polymerase B |  | Helicase / Helicase III (D5R-like helicase) |  | Lef-8 / Rbp2 |  | Lef-9 / Rbp1 |  | Lef-5 / TFIIIS-like transcription factor |  | P33 / Sulphydryl oxidase |  |
| --- | --- | --- | --- | --- | --- | --- | --- | --- | --- | --- | --- | --- |
|  | PSI | HMM | PSI | HMM | PSI | HMM | PSI | HMM | PSI | HMM | PSI | HMM |
| *Adelognathus sp.* | 0 | 0 | 0 | 0 | 0 | 0 | 0 | 0 | 0 | 0 | 0 | 0 |
| *Aleiodes sp.* | 0 | 0 | 0 | 0 | 0 | 0 | 0 | 0 | 0 | 0 | 0 | 0 |
| *Anoncus sp.* | 0 | 0 | 0 | 0 | 0 | 0 | 0 | 0 | 0 | 0 | 0 | 0 |
| *Dolichomitus sp.* | 1 | 1 | 1 | 0 | 2 | 2 | 1 | 0 | 0 | 0 | 0 | 0 |
| *Dusona sp.* | 0 | 0 | 0 | 0 | 0 | 1 | 1 | 1 | 0 | 0 | 0 | 0 |
| *Eumacrocentrus americanus* | 0 | 0 | 0 | 0 | 0 | 0 | 0 | 0 | 0 | 0 | 0 | 0 |
| *Lissonota sp.* | 0 | 0 | 0 | 0 | 0 | 0 | 0 | 0 | 0 | 0 | 0 | 0 |
| *Mesochorus sp.* | 0 | 0 | 0 | 0 | 0 | 0 | 0 | 0 | 0 | 0 | 0 | 0 |
| *Meterous sp.* | 0 | 0 | 0 | 0 | 0 | 0 | 0 | 0 | 0 | 0 | 0 | 0 |
| *Odontocolon sp.* | 0 | 0 | 0 | 0 | 0 | 0 | 0 | 0 | 0 | 0 | 0 | 0 |
| *Phanerotoma sp.* | 0 | 0 | 0 | 0 | 2+ | 2 | 1 | 1 | 0 | 0 | 1 | 1 |
| *Cotesia vestalis* | 2 | 1 | 1 | 1 | 2 | 1 | 2 | 1 | 0 | 0 | 0 | 0 |
| *Diachasma alloeum* | 0 | 0 | 0 | 0 | 0 | 0 | 0 | 0 | 0 | 0 | 0 | 0 |
| *Fopius arisanus* | 0 | 0 | 1 | 1 | 1 | 1 | 1 | 2* | 1 | 1 | 1 | 1 |
| *Macrocentrus cingulum* | 0 | 0 | 0 | 0 | 0 | 0 | 0 | 0 | 0 | 0 | 0 | 0 |
| *Microplitis demolitor* | 0 | 0 | 1 | 1 | 1 | 1 | 1 | 1 | 0 | 0 | 1 | 1 |
| *Venturia canescens* | 0 | 0 | 1 | 1 | 1 | 1 | 1 | 2* | 0 | 0 | 1 | 1 |

\*ORFs with hits are located next to each other on scaffolds and are fragmented due to pseudogenization
or intron presence
+Likely one gene fragmented across two contigs

In new wasp genomes, identification of at least one ancient core gene provided enough evidence to justify further exploration, while the absence of any hits led to the conclusion that specific wasp species are unlikely to produce virions or VLPs and exclude these species from further analysis. Proteins encoded by ancient core genes were identified in *Dolichomitus* (*DNApol, helicase, lef-8, lef-9*), *Dusona* (*lef-8, lef-9*), and *Phanerotoma* (*lef-8, lef-9*, *p33*).

Manual examination of protein alignments for each putative ancient core gene identified in wasp species (aligned with homologs from representative large DNA viruses and EVEs) confirmed protein identity and revealed that almost all hits were full length. Exceptions included one of the two *DNApol* genes from *C. vestalis* and one of the two *lef-9* genes from *Dolichomitus. Lef-8* sequences from *Phanerotoma* and *Dolichomitus* each had two ORFs matching this protein (N-terminal and C-terminal fragments). The *Phanerotoma* coding sequences lie on two different contigs, likely indicating broken assembly, and the *Dolichomitus* sequences lie near each other, suggesting that they are pseudogenized. The matches from *Dusona* were ORFs that were truncated compared to full length *lef-8* and *lef-9* sequences, and came from short scaffolds with low coverage, so were excluded from further analysis as likely contaminant sequences.

Phylogenetic reconstruction of ancient core gene trees revealed that genes of viral origin in each new wasp genome assembly came from a variety of sources (Figure 2, Supplementary Table 6). As expected, *Phanerotoma lef-8, lef-9*, and *p33* sequences grouped with the bracoviruses. *Dolichomitus DNApol, Helicase*, *lef-8*, and *lef-9* sequences all grouped with protein sequences from *Leptopilina boulardi* Filamentous Virus (LbFV), a large double-stranded DNA virus that is distantly related to hytrosaviruses and may represent a distinct virus family (Lepetit et al. 2017). The *C. vestalis* genome contains at least two ORFs of viral origin for *lef-8* and *lef-9* each. *C. vestalis* sequences fell into two groups: sequences clustered with bracoviruses (*helicase, lef-8*, and *lef-9*) or with LbFV (*DNApol, lef-8*, and *lef-9*). These data indicate that the sequenced *C. vestalis* samples contain two separate sources of viral genes; bracoviruses and a LbFV-like entity.

**Figure 2.**
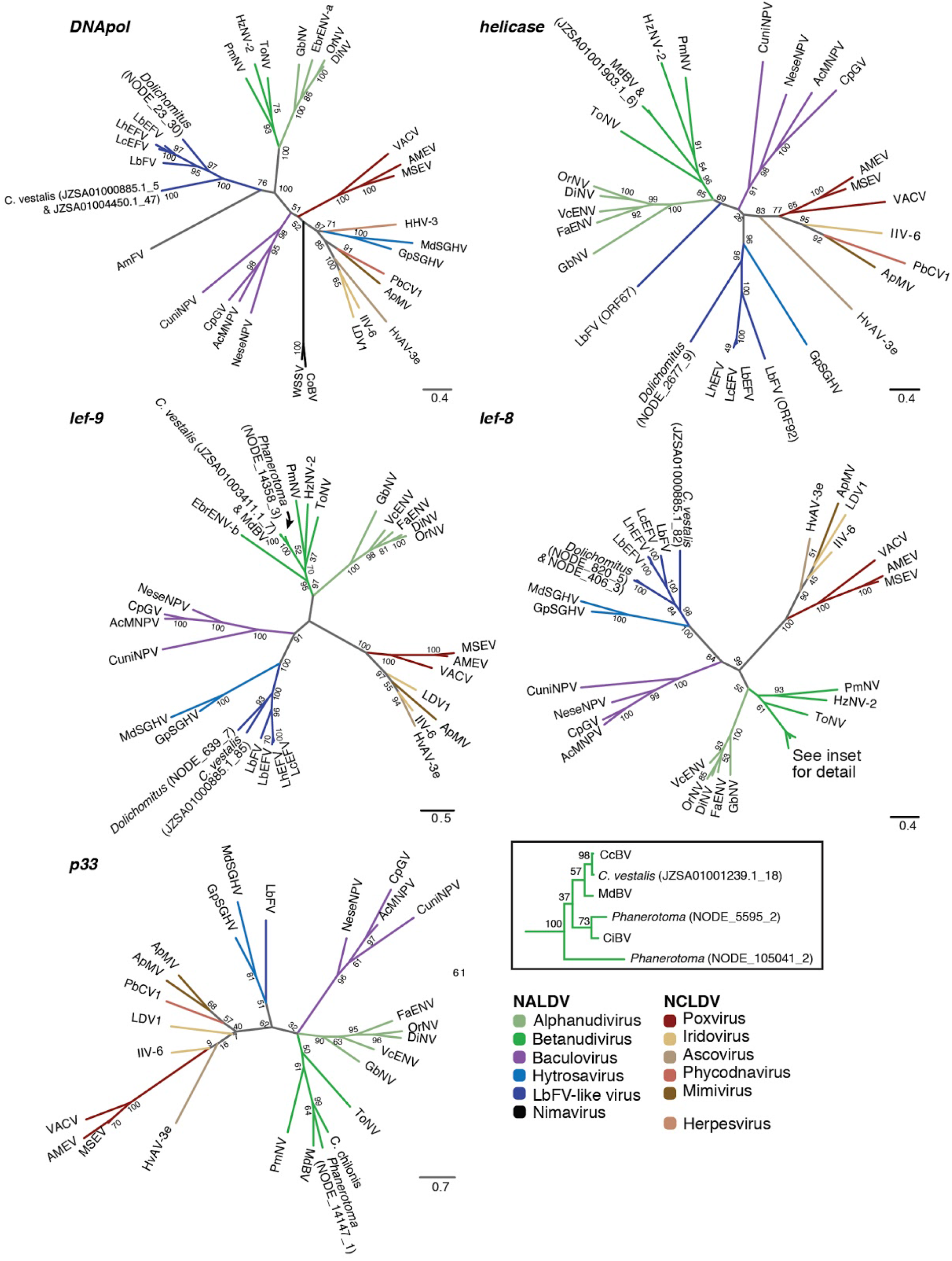
Genes of viral origin in wasp genomes are related to genes in large double-stranded DNA viruses. Maximum likelihood phylogenies constructed from protein sequences of individual genes are shown, with numbers on nodes indicating support from 100 bootstrap replicates. Colored branches indicate virus families or groups. The sequences used for alignment were obtained from *Microplitis demoltior* bracovirus (MdBV), *Cotesia congregata* bracovirus (CcBV), *Chelonus insularis* bracovirus (CiBV), a *C. chilonis* transcriptome, *Eurytoma brunniventris* endogenous nudivirus alpha (EbENV-a), *Eurytoma brunniventris* endogenous nudivirus beta (EbENV-b), *Tipula oleracea* nudivirus (ToNV), *Heliothis zea* nudivirus 2 (HzNV-2), *Penaeus monodon* nudivirus (PmNV), *Venturia canescens* endogenous nudivirus (VcENV), *Drosophila innubila* nudivirus (DiNV), *Oryctes rhinoceros* nudivirus (OrNV), *Fopius arisanus* endogenous nudivirus (FaENV), *Gryllus bimaculatus* nudivirus (GbNV), *Autographa californica* multiple nucleopolyhedrovirus (AcMNPV), *Cydia pomonella* granulovirus (CpGV), *Neodiprion sertifer* nucleopolyhedrovirus (NeseMNPV), *Culex nigripalpus* nucleopolyhedrosis virus (CuniNPV), *Glossina pallidipes* salivary gland hytrosavirus (GpSGHV), and *Musca domestica* salivary gland hytrosavirus (MdSGHV), *Leptopilina boulardi* filamentous virus (LbFV), *Leptopilina boulardi* endogenous filamentous virus (LbEFV), *Leptopilina heterotoma* endogenous filamentous virus (LhEFV), *Leptopilina clavipes* endogenous filamentous virus (LcEFV), *Apis mellifera* filamentous virus (AmFV), White spot syndrome virus (WSSV), *Chionoecetes opilio* bacilliform virus (CoBV), *Amsacta moorei* entomopoxvirus (AMEV), *Melanoplus sanguinipes* entomopoxvirus (MSEV), Vaccinia virus (VACV), Invertebrate iridescent virus 6 (IIV-6), LDV1, *Trichoplusia ni* ascovirus 3e (TnAV-3e), *Paramecium bursaria* chlorella virus 1 (PbCV1), *Acanthamoeba polyphaga* mimivirus (ApMV), Human herpesvirus 3 (HHV-3), Table S5. Sequences from the IVSPERs of the ichnovirus in *G. fumiferanae* and nimavirus *p33* homologs were not included because they were too divergent compared to the other included sequences. Scale bars indicate substitutions per amino acid residue. For ORFs found in wasp genomes, the accession number or scaffold name is followed by an underscore and the ORF number.

### Bracovirus nudivirus-like replication genes were detected in species in the microgastroid complex

As described above, bracoviruses were expected in the previously unannotated *C. vestalis* genome assembly and the new *Phanerotoma* genome assembly. These findings provide two proof-of-concept datasets to validate the approach for the identification of nudivirus-like replication genes beyond just the ancient core genes. Although genes from proviral segments are undoubtedly present in the datasets, their annotation was not attempted due to the fragmented nature of the genome assemblies and the lack of conserved genes that could be used as a proviral segment diagnostic. The first strategy used for the identification of nudivirus-like replication genes involved alignment of ORFs from each genome assembly to the custom diamond database. The second strategy was designed to identify an expanded set of nudivirus-like EVEs in each genome assembly, and involved adding nudivirus-like replication genes from *M. demolitor, C. congregata*, and *C. inanitus* to the custom diamond database and repeating the diamond search. The existing nudivirus-like replication gene models from bracoviruses are the products of careful manual annotation of sequence data. These manually curated sequences are more likely to generate database search hits compared to searches against the set of insect virus genomes available in public databases because they are less divergent than other nudiviruses; these genes are the products of a single integration event and share a common ancestor ~100mya (Murphy et al. 2008).

Using the first search strategy, 18 ORFs similar to nudivirus and baculovirus genes were identified on 18 contigs in the *Phanerotoma* assembly ranging in size from 885bp to 12,791bp. The GC content of these contigs was not significantly different from BUSCO-containing contigs, but on average their coverage was slightly higher (Supplementary Figure 2, 28x compared to 9x, *p* < 0.05). The second strategy for identification of nudivirus-derived genes found 13 of the original 18 ORFs plus 37 additional genes (55 total, Supplementary Table 7). A search of all ORFs in the *C. vestalis* genome against the custom diamond database revealed 29 ORFs matching to nudivirus or baculovirus coding sequences located on 24 contigs. The GC content of contigs containing nudivirus-like genes was not significantly different than BUSCO-containing contigs (mean 29.7% compared to 29.2%, respectively, *p* = 0.30). It was not possible to create scaffold coverage and GC content plots for *C. vestalis*, because the sequencing reads used to make the genome assembly were not available in the NCBI Sequence Read Archive. A broader search using the second strategy as described above identified 13 of the original 29 ORFs, plus an additional 78 hits (106 total, Supplementary Table 7).

To assess the completeness of the set of bracovirus replication genes identified in each assembly, a tally of the genes conserved in bracoviruses and also nudiviruses that are presumed to be essential for virion production was conducted. The *M. demolitor* genome contains at least one copy of 19 of the 33 nudivirus core genes (18 of these are also core genes in baculoviruses, Burke 2019). All of the 19 genes conserved in nudiviruses and *M. demolitor* were present in the set of nudivirus-like genes identified in *Phanerotoma*, and only one gene, *pif-4*, was missing in the *C. vestalis* assembly. These results indicate that the methods used were appropriate for the identification of the presence of EVEs that are likely to produce bracoviruses. However, the genome assemblies were sufficiently fragmented that there was not enough information to confidently assess the placement of genes or the presence of synteny between nudivirus-like genes in microgastroid species. The fragmented nature of the new genome assemblies and the coarse nature of the methods used made it possible that a complete list of EVE genes in the *Phanerotoma* and *C. vestalis* genomes was not fully identified. A more detailed and comprehensive analysis will be appropriate if and when assemblies with larger scaffold sizes are available for these species.

### Ichnovirus Structural Protein Encoding Regions (IVSPERs) were detectable but not always present where expected

Given that no ancient core genes have been documented as consistently present in ichnovirus-producing wasp genomes to date, the strategy used for the identification of IVSPER genes in wasp genome assemblies relied upon homology to the IVSPERs from *H. didymator* and *G. fumiferanae*. IVSPER genes were identified in the *Lissonota* genome assembly associated with ten scaffolds ranging in size from 1,227bp to 44,485bp (Supplementary Figure 2). The protein sequence percentage identity to *G. fumiferanae* IVSPER genes from diamond searches was 28-86%, with an average of 57%. Like the scaffolds containing EVEs in *F. arisanus* and *M. demolitor*, in *Lissonota* the IVSPERs were located on scaffolds that did not significantly differ in coverage from scaffolds that contain BUSCO genes (Supplementary Figure 2). This indicates that as in other IV-producing wasps, the IVSPER genes are integrated into the genome of *Lissonota* (Béliveau et al. 2015; Volkoff et al. 2010). Scaffolds containing IVSPERs had a slightly higher GC content (39.7%) compared to BUSCO scaffolds (37.8%, *p*<0.05). *Lissonota* IVSPERs had high levels of synteny with IVSPERs previously identified in the banchine *G. fumiferanae*, indicating that the viral integration event occurred in a shared ancestor of these wasp species (Figure 3). Twenty-four genes were shared between these two species and were also present in the IVSPERs in *H. didymator*, while another 25 were present in the two banchine species only. The four genes linking IVs to NCLDVs (*ssp1, DNApol, D5 primase*, and *helicase*) were present in *Lissonota* as well as *G. fumiferanae*, where they were discovered.

**Figure 3.**
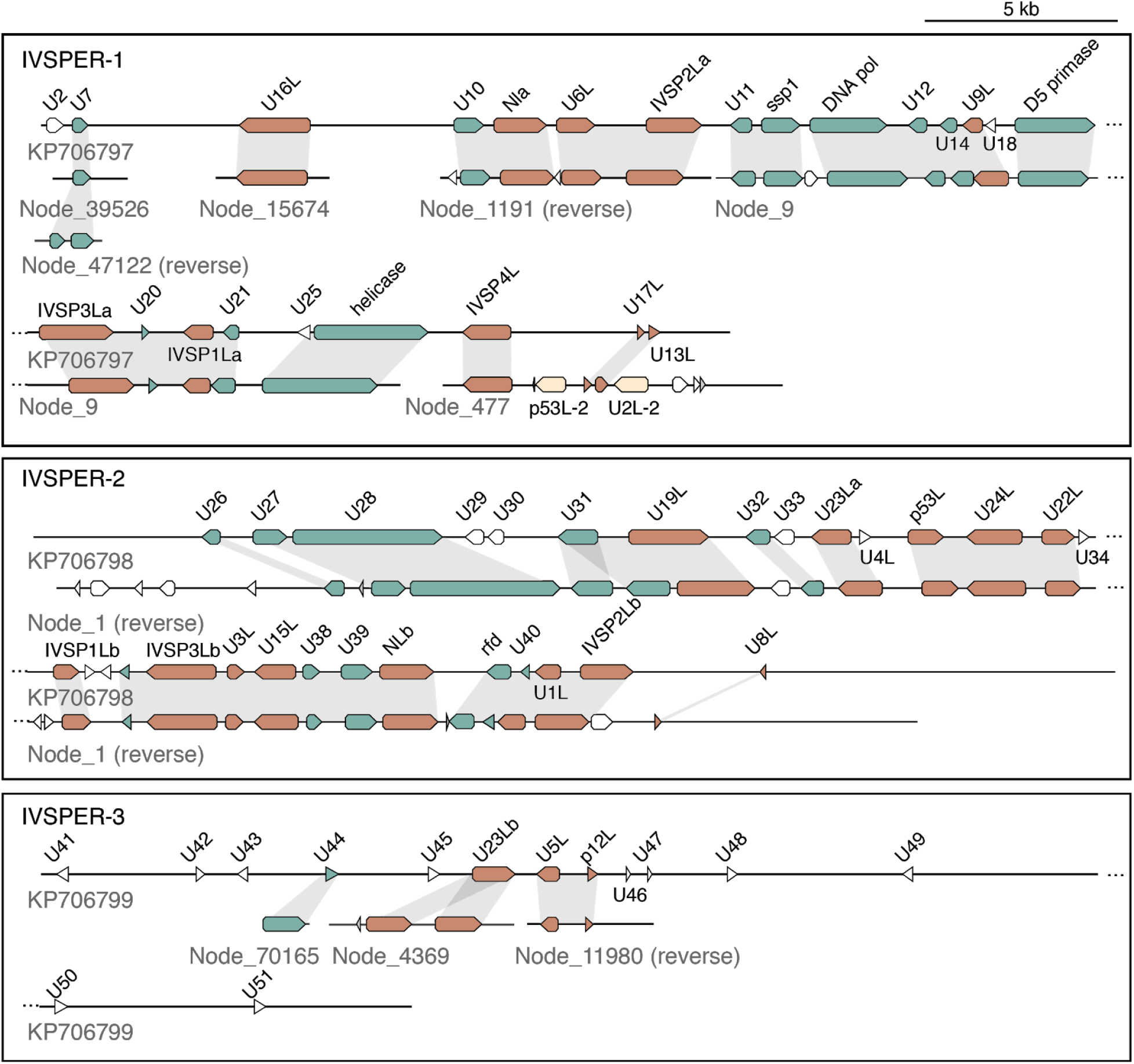
IVSPER genes identified in the *Lissonota* genome and synteny with *G. fumiferanae. G. fumiferanae* sequences are shown above with *Lissonota* sequences shown below. Homologous genes with synteny between the two species are indicated by grey shading. Some genes are present in banchine and campoplegine species (*G. fumiferanae, Lissonota*, and *H. didymator*; colored orange) while others are present only in the two banchine species presented here (blue). Genes colored beige are homologous to other IVSPER genes but are not detected in syntenous regions of the *G. fumiferanae* genome.

There were no significant hits to any ORFs from the remaining ten new genome assemblies or the previously published genome assemblies. The only exception was a match to a hypothetical protein (U44, AKD28091.1) in the *V. canescens, M. cingulum, F arisanus* and *Lissonota* ORFs. BLAST of the *G. fumiferanae* U44 revealed that this protein has strong similarity to other hymenopteran proteins, making it unclear whether the ORFs identified are viral in origin, or merely wasp genes. The absence of hits to IVSPER genes in *Dusona* is notable because this species belongs to the Campopleginae, where all taxa were assumed to have ichnoviruses. The absence of IVSPERs and related genes in *Mesochorus* (Mesochorinae) and *Anoncus* (Ctenoplematinae) is also notable as they share a common ancestor with the Campopleginae and Banchinae. In addition, the absence of IVSPERs in *Dusona* suggest that IVs might actually be limited to a subset of species within the campoplegine wasps.

### Novel filamentous virus-like genes were identified in the *C. vestalis* and *Dolichomitus* genomes

The identification of several ancient core genes related to LbFV genes in the *C. vestalis* and *Phanerotoma* genomes provided a strong hint that these species contain EVEs or are “contaminated” with sequences from an actively replicating viral infection. This result motivated expansion of the search for LbFV-like genes beyond the ancient core genes initially identified to look for signatures of wasp genome integration. The genome of *C. vestalis* was available on NCBI in an unannotated state because the assembly is somewhat fragmented. In addition to genes originating from bracoviruses, HMM and PSI-BLAST searches identified four additional ancient core genes in the *C. vestalis* genome (two copies of *DNApol*; *lef-8* and *lef-9*).

In order to effectively identify LbFV-like genes in wasp genomes, better annotation of genes in LbFV was attempted. Previous studies identified homologs for *DNApol, helicase, lef-8, lef-9, pif-0 (p74), pif-2, pif-5, ac81*, and *odv-e66* in LbFV (Di Giovanni et al. 2020; Lepetit et al. 2017; Kawato et al. 2018). Using HMMs built with homologs from other insect-infecting large dsDNA viruses, it was possible to identify the following additional genes in LbFV: *lef-4* (ORF107)*, 38K* (ORF19), and *pif-1* (ORF32) (Supplementary Table 8). In addition to the four LbFV-like ancient core genes, the results from diamond and HMM searches revealed nine more LbFV-like genes in the *C. vestalis* genome. Genes similar to LbFV genes were identified on three *C. vestalis* genomic scaffolds (Figure 4, accession doi: 10.15482/USDA.ADC/1504545). *DNApol, lef-4, lef-8, lef-9* and 25 other neighboring genes encoding hypothetical proteins with typical viral architecture (short, closely spaced, no introns) are located on the first scaffold (JZSA01000885.1) next to four eukaryotic genes encoding uncharacterized proteins: two with similarity to hymenopteran species and another with similarity to *Helicoverpa armigera*. The second scaffold (JZSA01004450.1) contains a partial (50% full length) copy of a viral DNApol gene situated among seven genes with eukaryotic architecture and strong similarity to other hymenopteran genes, including *syntaxin-7, adenylate cyclase type 2, importin-4*, and *forkhead box-like* genes (Figure 4). The final scaffold (JZSA01007324.1) contains *lef-3, 38K, p74, pif-2, pif-5, ac81, odv-e66, putative lecithin:cholesterol acyltransferase* genes and 52 other genes with viral architecture. Interspersed are two genes that have eukaryotic architecture with significant similarity to genes encoding uncharacterized proteins in other hymenopteran species (Figure 4). Alignment of sequence expression data provided evidence that 6 of the 13 eukaryotic genes are expressed and thus represent high-confidence annotations.

**Figure 4.**
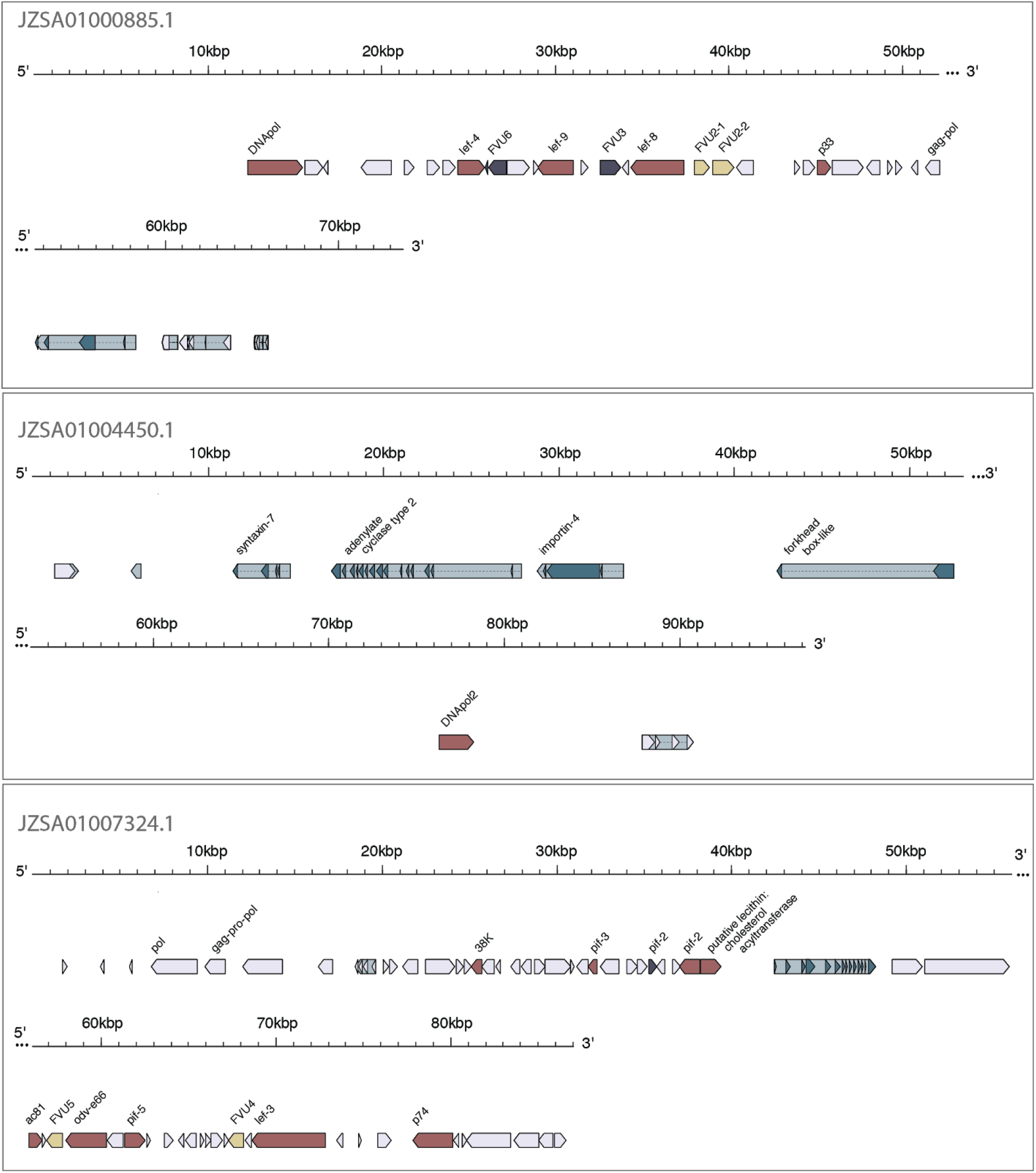
Regions of the *C. vestalis* genome similar to the *Leptopilina boulardi* Filamentous Virus (LbFV). Genes with homology to LbFV are highlighted red, pseudogenes as a deep purple. Genes with homology to LbFV-like genes in *Dolichomitus* are shaded yellow or purple if pseudogenized (FVU = Filamentous Virus Unknown protein encoding gene). Other predicted genes are shaded in grey. Genes with blue background shading have eukaryotic structure (introns and exons), and of these, genes with dark blue colored exons are “high confidence” given evidence from EST alignments.

The overall organization of genes in these loci suggest that LbFV-like genes are present in the wasp genome in three different locations; two likely representing primary integration events constituting large portions of the LbFV-like ancestor’s genome, and the third possibly representing a secondary integration event in which part of the viral *DNApol* gene was transferred to a different region of the wasp genome. For both of the putative primary integration events, the ends of strings of genes of with viral architecture are associated with remnants of retroviral genes such as *pol, gag-pol*, and *gag-pro-pol*, suggesting retroelements as a possible source for entry of viral genes into the wasp genome (Desjardins et al. 2008).

Two genome assemblies are available for *C. vestalis* on NCBI. The two assemblies were generated from wasp populations located in Andong, South Korea (ASM95615v1) and Hangzhou, China (ASM167554v1). Full length versions of all of the genes of viral origin were present in both assemblies (except *ac81*, which was only present at 80% length in the Hangzhou assembly, located at the end of a scaffold) with nucleotide sequence identity ranging from 97-99%. The presence of slightly divergent genes of viral origin in both assemblies further suggests that these genes are integrated into the wasp genome for both populations of *C. vestalis*.

A total of 18 contigs in the *Dolichomitus* genome assembly (ranging in size from 312bp to 39.5kb) contained genes related to LbFV. These contigs were too short in length to determine whether any genes with eukaryotic architecture were flanking the virus-derived genes. The GC content of these contigs was significantly lower than BUSCO-containing contigs (mean 37.4% compared to 39.6%, *p* <10^−4^), and on average their coverage was higher (28x compared to 7x, *p* < 10^−9^ on log_10_ transformed values, Supplementary Figure 2). After manual annotation of these contigs, diamond analysis revealed that 24 of the total 149 ORFs had similarity to LbFV genes, including *DNApol, lef-3, lef-4, lef-5, lef-8, lef-9, pif-0, pif-1, pif-2, pif-3, pif-5, 38K, helicase, helicase2*, and *ac81* (Figure 5). Although the cumulative size of the “viral” contigs was 199kb, the non-degenerate size was approximately 114kb because many contigs were repetitive, with up to three divergent copies of distinct genes present among the set (Figure 5). BLAST alignment identity for LbFV genes to *Dolichomitus* ORFs ranged from 22.9-42.7% (mean 31.2%, median 32%), while identity among paralogous *Dolichomitus* ORFs was 30.4-95.83% (average 62.8%, median 62.9%). Comparison of the *Dolichomitus* and *C. vestalis* genes with viral architecture flanking LbFV-like genes revealed homologs of six genes that are common to the two wasp genome assemblies but are not present in LbFV, and were given the gene names Filamentous Virus Unknown (FVU) 1 through 6 (Figure 5). The divergence of viral genes in *Dolichomitus* compared to LbFV often made it difficult to determine whether an ORF represents a full-length gene. However, homology between ORFs on different contigs in *Dolichomitus* revealed that some ORFs represent fragments of pseudogenized genes (Figure 5). Of all of the ORFs predicted on the 18 *Dolichomitus* contigs, 33 are fragments of a total of 14 pseudogenized genes (ranging from 1 to 6 per contig). On three contigs, core genes that are considered essential for replication were inactivated: *38K* on Node_1, *p74* and *pif-5* on Node_10, *lef-4* on Node_71 and *lef-8* on NODE_820. The presence of degenerated essential genes provides strong evidence that at least these four LbFV-like contigs are endogenous in the *Dolichomitus* genome. These contigs are not likely to be part of an archetypal virus genome actively replicating in *Dolichomitus*, because the products of these genes would be non-functional or absent and are essential for virion formation. The presence of several syntenous copies of viral genes could be the product of multiple integration or genome locus duplication events that have substantially diverged over time. If all of the LbFV-like contigs were endogenous in the *Dolichomitus* genome, it is noted that at least one copy of each LbFV-like gene with essential functions was intact across the contigs cumulatively. Thus, it is theoretically possible that this *Dolichomitus* wasp produced virus-like particles by assembling protein products from intact LbFV-like genes that are dispersed throughout the *Dolichomitus* genome.

**Figure 5.**
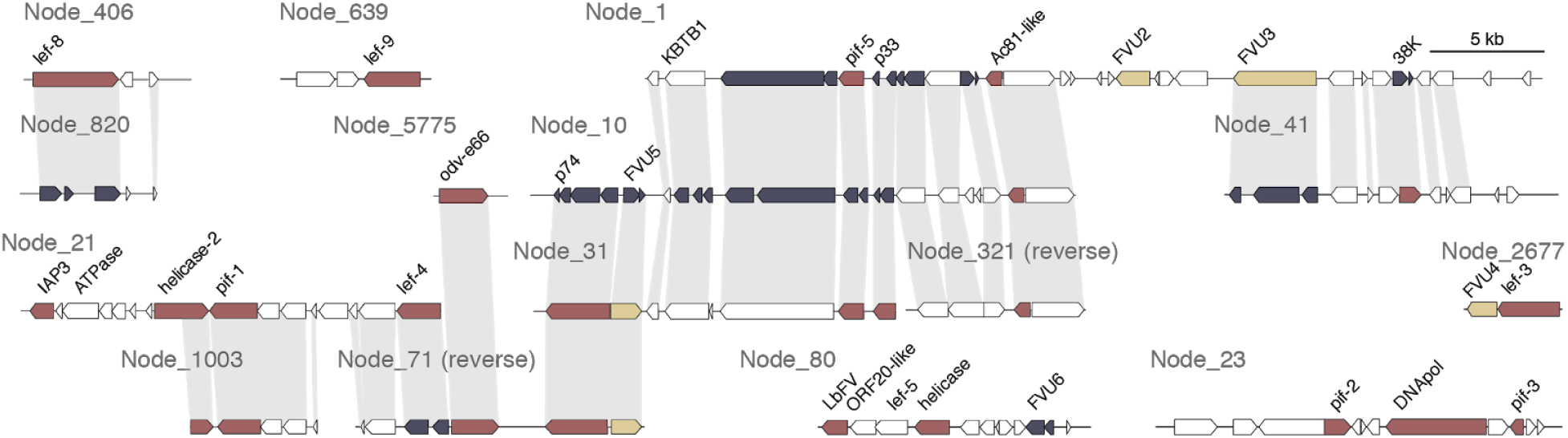
Regions of the *Dolichomitus* genome similar to the *Leptopilina boulardi* Filamentous Virus (LbFV). Homologous genes with synteny between contigs are indicated by grey shading. Genes with homology to LbFV are highlighted red, genes with homology to LbFV-like genes in *C. vestalis* are shaded yellow, and other predicted genes are colored white. FVU = Filamentous Virus Unknown protein encoding gene. ORFs shown in deep purple represent pieces of pseudogenes that can be found intact on other contigs. Three short nodes with incomplete gene sequences were not included in this figure (NODE_51377 and NODE_267127 containing *pif-0*, and NODE_236835 containing *lef-9*).

### EVEs in parasitoid wasp species are distantly related to other arthropod DNA viruses

After annotation of the genes of viral origin in all of the new wasp genome assemblies, a list of genes emerged as conserved in many if not all viruses and EVEs derived from within the NALDVs. In addition to the ancient core genes (*DNApol, helicase, lef-5, lef-8, lef-9*, and *p33*), the recent description of nimavirus EVEs present in crustacean genomes identified *pif-0, pif-1, pif-2, pif-3*, and *pif5* as core genes present in NALDVs (Kawato et al. 2018). Based upon new data from EVEs related to LbFV, *ac81* can be added to a core gene set for viruses that infect insects (Supplementary Table 8). Protein sequences from these 12 genes were used to construct a multi-gene phylogeny and examine the relationships between viruses and EVEs (Figure 6).

**Figure 6.**
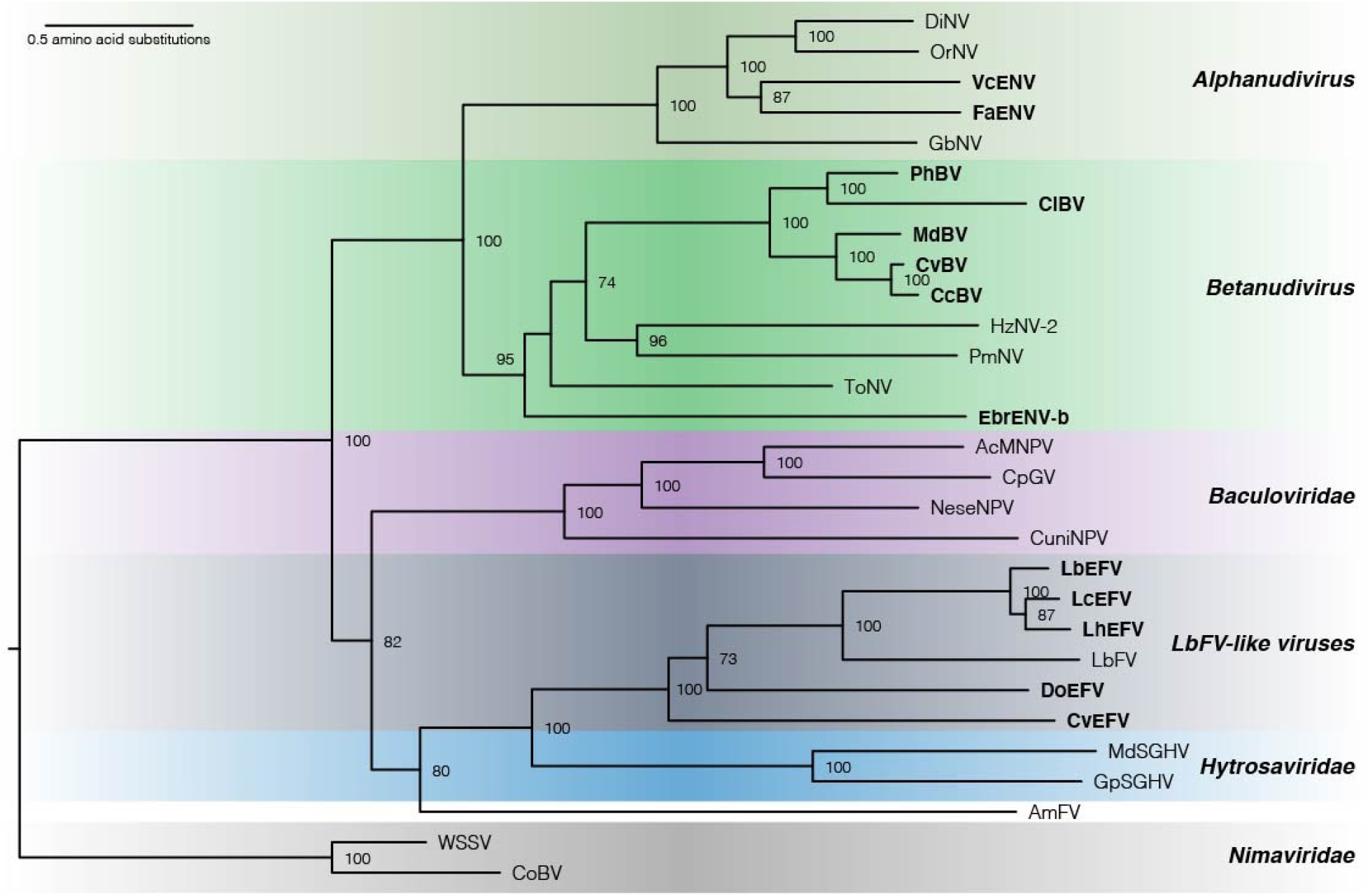
Phylogenetic analysis of arthropod infecting large double-stranded DNA viruses and parasitoid EVEs. Relationships were derived using a maximum likelihood analysis from 12 core genes with a total of 5003 characters from concatenated amino acid sequences. Bootstrap values over 50% are indicated near the relevant node. Scale bar indicates average number of amino acid substitutions per site. Names of EVEs are highlighted with bold type. Taxa are named as in Figure 1, with the addition of *Phanerotoma* bracovirus (PhBV), *Cotesia vestalis* bracovirus (CvBV), *Dolichomitus* endogenous filamentous virus (DoEFV), and *Cotesia vestalis* endogenous filamentous virus (CvEFV). EbrENV-a was omitted because only one sequence (DNA polymerase) was available.

As expected, bracoviruses from the Cheloninae (*Phanerotoma* and *C. inanitus*) were most closely related to each other, as were bracoviruses from both wasps belonging to *Cotesia* within the Microgastrinae (*C. congregata* and *C. vestalis*) (Figure 6). The EVEs related to LbFV are referred to as *Dolichomitus* Endogenous Filamentous Virus (DoEFV) and *C. vestalis* Endogenous Filamentous Virus CvEFV). These EVEs are as divergent from each other as they are from LbFV. Despite their names, the filamentous viruses from *L. boulardi* and *A. mellifera* are separated on the phylogeny by hytrosaviruses and do not appear to belong to the same virus family (Figure 6).

## Discussion

This study investigated the presence and distribution of EVEs in one of the most astonishing radiations on Earth (parasitoid wasps from the Ichneumonoidea) and hints at the massive untapped number and diversity of viral acquisition events. As of early 2019 when this study began, only five genomes from the Ichneumonoidea were available in the National Center for Biotechnology Information database, but with advances in sequencing technologies, rapid growth of this number is expected. The availability of genome sequence data from 11 new species representing diverse lineages from within the Ichneumonoidea provided a unique opportunity to identify genes of viral origin. The identification of sets of genes involved in viral replication is suggestive of the production of virions or VLPs that have a significant impact upon parasitoid wasp biology. Further, characterizing the presence or absence of EVEs in relatives of wasp species known to produce virions or VLPs provides information about the origination of these features and their species distribution.

A new approach was needed in order to screen wasp genomes for genes of viral origin in a consistent and high-throughput manner. The approaches designed here were validated using six previously published genomes from the Ichneumonoidea with known presence or absence of EVEs. The first, simplest approach used to identify genes of viral origin was to screen all ORFs against a streamlined form of the NCBI nr database. However, this screen resulted in false positives due to the presence of many hits to viral genes that are likely to be frequently transferred between eukaryotic and viral genomes. The second approach therefore focused upon identification of ancient core genes, or genes that serve essential functions in the production of virions in dsDNA viruses. With this approach, it was possible to verify the presence of bracoviruses in the previously published genomes of *M. demolitor* and *C. vestalis*. The presence of genes derived from nudivirus acquisition events was also recovered with this method in *F. arisanus* and *V. canescens*. The ancient core gene approach also accurately showed that there are no virus gene acquisitions in the *D. alloeum* or *M. cingulum* genomes.

Having established a method that could pinpoint which species to focus upon further, the new genome assemblies were searched for the presence of ancient core genes. If ancient core genes were identified, their evolutionary history was reconstructed with phylogenetic trees with representatives of the diversity of large dsDNA viruses. Ancient core genes are not necessarily monophyletic, in fact, many seem to be acquired from eukaryotic replication systems independently in NALDVs and NCLDVs (Iyer et al. 2004, 2006, 2003; Kool et al. 1994). Thus, the weak support at basal nodes of the phylogenetic trees in Figure 2 was expected, and rather than attempting to elucidate relationships between ancient divergence events in viral evolution, these trees functioned to categorize viral endogenization events into groups with viral ancestors.

As expected, a bracovirus was identified in the microgastrine braconid, *Phanerotoma*, and an ichnovirus in the banchine ichneumonid, *Lissonota*. The screen for ancient core genes then identified two major findings that were not expected *a priori*: 1) the presence of LbFV-related genes in *C. vestalis* (Braconidae: Microgastrinae) and *Dolichomitus* (Ichneumonidae: Pimplinae) genomes, and 2) the absence of an ichnovirus in the campoplegine ichneumonid, *Dusona*. The preceding results will be discussed further below.

### Both nudiviruses and LbFV-like viruses are common endogenous virus progenitors

Prior to this study, endogenous viruses discovered in parasitoid wasp species revealed that virus co-option from nudivirus ancestors has occurred independently on at least four occasions (Bezier et al. 2009, Burke et al. 2018, Pichon et al. 2015, Zhang et al. 2020), all in koinobiont endoparasitic wasps. Out of all of the major groups of insect-infecting viruses, the predominance of nudivirus ancestors of EVEs suggested specific, as yet unknown conserved features of nudiviruses may predispose this group to integration and establishment in wasp genomes (Strand & Burke 2020, 2014). Recently, the first endogenization event from an ancestor other than nudiviruses and the ichnovirus ancestor was identified in *Leptopilina boulardi* (Figitidae), a koinobiont endoparasitoid wasp that produces VLPs or Mixed Strategy Endocytic Vesicles (MSEVs) (Di Giovanni et al. 2020; Heavner et al. 2017; Rizki & Rizki 1990). The ancestor of the EVE in *L. boulardi* (henceforth referred to as LbEFV) was a behavior-manipulating filamentous virus that also infects *L. boulardi* (LbFV), although the two are relatively distantly related (Di Giovanni et al. 2020). This virus was only recently identified and although it clearly belongs to the NALDVs, it is so divergent from other virus families that it most likely represents the first discovered member of a new virus family (Lepetit et al. 2017).

This study identified two more independently-derived instances of endogenization of a LbFV-like entity into parasitoid wasp genomes, making LbFV-like viruses a second group of viruses that seem to be predisposed to endogenization. Both *C. vestalis* and *Dolichomitus* possess intact versions of many of the genes necessary to make virions or VLPs, justifying future investigation of the reproductive secretions of these wasp species. The LbFV-like virus in *Dolichomitus* also represents the first endogenized virus in an idiobiont ectoparasitoid, suggesting that this virus may function differently for parasitism success. Additionally, given that *C. vestalis* already produces a bracovirus, investigation of the role of LbFV-like genes in this species could address whether LbFV-like virions are also produced and how both types of virus might impact parasitism success and/or behavior. It is noted that non-bracovirus filamentous particles have been observed in *Cotesia* species previously. Filamentous virions with similar morphology have been observed in *C. congregata* and *C. marginiventris* (as well as the ichneumonid *Eriborus terebrans* (formerly *Diagema terebrans*) (Styer et al. 1987; Hamm et al. 1990; Krell 1987; Townes 1965). Although no genetic data are available for the filamentous viruses in other *Cotesia* species, these viruses and LbFV have similar morphogenesis (capsid production in cell nuclei, acquisition of an envelope in the cytoplasm) and may be derived from the same type of viral ancestor (Varaldi et al. 2006; Buron et al. 1992).

### “Ichnoviruses” represent two independent, relatively recent virus acquisition events

The genus “*Ichnovirus*” was coined to describe a type of virus that replicates in wasp ovaries and is injected into host insects during oviposition (Lefkowitz et al. 2018). This type of virus had a lenticular capsid surrounded by two unit membrane envelopes and a polydisperse genome including multiple double-stranded, circular DNAs of different sizes and coding capacities (Stoltz et al. 1984). At that time, it was not known that ichnoviruses were actually heritable endogenous entities, unlike archetypal viruses. It is more appropriate to group and name EVEs according to their independent evolutionary origins, which requires knowledge of the evolutionary history of wasp species that possess EVEs. The hyperdiversity of species within the Ichneumonidae has made their evolutionary relationships historically difficult to resolve. However, recent studies (Quicke et al. 2009; Klopfstein et al. 2019; Bennett et al. 2019) have confirmed results from a single gene phylogeny to show that Campopleginae and Banchinae are not closely related to each other. Two previous studies have described the IVSPERs in a single member of each of these subfamilies, and from extensive overlap of the catalogue of IVSPER genes, concluded that these EVEs came from similar, if not identical virus ancestors (Béliveau et al. 2015; Volkoff et al. 2010). Beliveau et al (2015) suggested two evolutionary scenarios giving rise to the presence of IVSPERs in these two divergent wasp subfamilies. First, that there was a single IV ancestor, in which a NCLDV integrated into the genome of an ancestor of banchine and campoplegine wasps. If no IVs are observed in wasp species belonging to the subfamilies that also descend from the common ancestor of banchine and campoplegine wasps, it implies that the capacity to produce IVs was lost in all of these species, although traces of IVSPER genes could remain in their genomes. The second scenario involves two separate integration events involving very similar viral ancestors (related to NCLDVs) that took place separately in the ancestors of banchine and campoplegine wasps.

Our data support the second scenario involving two independent integration events in campoplegine and banchine wasp species. First, based upon the phylogeny generated by Sharanowski et al. (2020), two of the wasp species with new genome assemblies, *Anoncus* (Ctenopelmatinae) and *Mesochorus* (Mesochorinae), are “intervening taxa”, *ie*. they do not belong to either of the IV-producing subfamilies but descend from the common ancestor of these subfamilies. Both of these genomes lack ancient viral core genes, common dsDNA virus replication genes, and IVSPER-like genes, intact or otherwise. Second, the unexpected absence of IVSPERs in *Dusona* suggests that ichnoviruses are not present in all campoplegine species, and may be limited to a clade containing *Hyposoter, Tranosema, Campoletis*, and *Casinaria* while excluding *Dusona* and possibly *Campoplex* (Sharanowski et al. 2020, (Volkoff et al. 2010; Pichon et al. 2015). Determining the distribution of EVEs among species is important because it will help to inform whether other viral acquisition events (such as the nudivirus in the campoplegine *V. canescens*) are truly replacement events or represent acquisition into a wasp ancestor that ancestrally lacked an EVE. Third, the architecture of the IVSPERs within wasp genomes also provides evidence of recent acquisition. The IVSPERs in *H. didymator*, *G. fumiferanae* and now *Lissonota* (all Banchines) are each located in three compact clusters within wasp genomes, respectively. This level of clustering is similar to the architecture observed for the recently-derived endogenous nudivirus in *F. arisanus* (nine major clusters), and lies in contrast to the extensive spread of nudivirus-like genes in the *M. demolitor* genome representative of the bracovirus-producing microgastrine wasps (Burke et al. 2014; Burke, Simmonds, et al. 2018; Burke, Walden, et al. 2018). As the spread of virus-derived wasp genes throughout wasp genomes is thought to be a relatively neutral process that occurs over evolutionary time (Burke et al. 2014), the limited spread of the IVSPERs in wasp genomes indicates that their integration occurred relatively recently. Finally, there are morphological differences between the campoplegine and banchine ichnoviruses that also suggest their independent origins (Béliveau et al. 2015). While a close relative of the “ichnoviruses” has yet to be discovered in a non-endogenous form, it is possible that the ichnovirus ancestor is also a common progenitor of endogenous associations in parasitoid wasps.

## Concluding remarks

This study has shown that sequencing parasitoid wasp genomes can reveal novel instances of EVE acquisitions in the Ichneumonoidea. The generation of new genomic data from just eleven species and re-analysis of six publicly available genomes has identified two independently-derived, novel EVEs. Additionally, the new genomes generated sequence data for two more EVEs that were expected in two species based upon their clade membership. Following the proposed use of a standardized nomenclature toward binomial species names for viruses (ICTV) (Siddell et al. 2020), revision of the genera *Bracovirus* and *Ichnovirus* and the family Polydnaviridae should be considered to reflect their endogenous nature and their evolutionary history. Additionally, coining a name for the LbFV-like viruses would be useful given their prevalence as EVEs in parasitoid wasps. Overall, the data presented in this study hints that the diversity of viral co-option events extends much further and is more common than just two rare, ancient events previously referred to as bracoviruses and ichnoviruses. Rather, the acquisition of EVEs from different groups of viral pathogens creates an ample source of variation for the recurrent evolution of diverse parasitism-based virulence strategies in parasitoid wasps.

## Supporting information

Supplementary Tables 4 through 8

## Acknowledgements

The authors would like to acknowledge Michael Strand for feedback upon the manuscript. This work was supported by the US National Science Foundation (DEB-1622986 to G.R.B. and DEB-1916788 to G.R.B. and B.J.S.), the USDA National Institute of Food and Agriculture Hatch project (1013423 to G.R.B.), and the Natural Sciences and Engineering Research Council (NSERC) Discovery Program (B.J.S.).

## Data availability

The data underlying this article are available in the National Center for Biotechnology Information at https://www.ncbi.nlm.nih.gov, and the Ag Data Commons at https://data.nal.usda.gov, and can be accessed with accession numbers given throughout the manuscript, figures, tables, and supplementary information.

