## Supplementary tables and figures for "Endogenization from diverse viral ancestors is common and widespread in parasitoid wasps"

Supplementary Table 1. Voucher information for species subjected to genome sequencing.

| **Voucher Code** | **Assembly accession** | **Genus species** |
|---|---|---|
| PSUC_FEM 10030000_I15696 | JABSMQ000000000 | *Anoncus sp.* |
| PSUC_FEM 10030001_I15694 | JABFVB000000000 | *Adelognathus sp.* |
| PSUC_FEM 10030002_I15713 | JABMBT000000000 | *Odontocolon sp.* |
| PSUC_FEM 10030003_I15695 | JABSMS000000000 | *Aleiodes sp.* |
| PSUC_FEM 10030004_I15704 | JABSMT000000000 | *Eumacrocentrus americanus* |
| PSUC_FEM 10030005_I15702 | JAAXZA000000000 | *Dolichomitus sp.* |
| PSUC_FEM 10030008_I15710 | JABMBR000000000 | *Meteorus sp.* |
| PSUC_FEM 10030009_I15714 | JABMBU000000000 | *Phanerotoma sp.* |
| PSUC_FEM 10030011_I15709 | JABMBV000000000 | *Mesochorus sp.* |
| PSUC_FEM 10030012_I15708 | JABMBS000000000 | *Lissonota sp.* |
| PSUC_FEM 10030013_I15703 | JABSMR000000000 | *Dusona sp.* |

Supplementary Table 2. Read data and assembly statistics for parasitoid wasp genome assemblies

| **Species** | **Raw reads sequenced (total nt)** | **Processed reads (total nt)** | **Number of genome scaffolds or contigs (total nt)** | **N50 scaffold size** | **Mean coverage** |
|---|---|---|---|---|---|
| *Adelognathus sp.* | 58.9 million (5.9 Gb) | 31.7 million (3.5 Gb) | 202,891 (267 Mb) | 2,095 nt | 11x |
| *Aleiodes sp.* | 82.8 million (8.3 Gb) | 46.0 million (5.2 Gb) | 72,851 (181 Mb) | 6,330 nt | 35x |
| *Anoncus sp.* | 61.4 million (6.1 Gb) | 33.2 million (3.7 Gb) | 332,752 (226 Mb) | 1,110 nt | 12x |
| *Dolichomitus sp.* | 65.4 million (6.5 Gb) | 35.8 million (4.0 Gb) | 294,254 (244 Mb) | 1,083 nt | 14x |
| *Dusona sp.* | 64.9 million (6.5 Gb) | 36.2 million (4.0 Gb) | 107,841 (234 Mb) | 3,885 nt | 29x |
| *Eumacrocentrus americanus* | 63.7 million (6.4 Gb) | 35.1 million (3.9 Gb) | 55,733 (117 Mb) | 3,852 nt | 35x |
| *Lissonota sp.* | 77.2 million (7.7 Gb) | 43.4 million (5.0 Gb) | 187,656 (233 Mb) | 2,166 nt | 24x |
| *Mesochorus sp.* | 88.7 million (8.9 Gb) | 49.5 million (5.6 Gb) | 85,631 (195 Mb) | 3,953 nt | 30x |
| *Meteorus sp.* | 75.0 million (7.5 Gb) | 42.4 million (4.9 Gb) | 13,843 (123 Mb) | 31,843 nt | 54x |
| *Odontocolon sp.* | 65.9 million (6.6 Gb) | 35.4 million (3.9 Gb) | 101,654 (173 Mb) | 3,208 nt | 25x |
| *Phanerotoma sp.* | 85.9 million (8.6 Gb) | 48.0 million (5.6 Gb) | 423,888 (324 Mb) | 1,222 nt | 13x |
| **Previously published** | | | | | |
| *Cotesia vestalis* | | | 9,156 (186 Mb) | 46,055 nt | |
| *Diachasma alloeum* | | | 3,968 (389 Mb) | 645,483 nt | |
| *Fopius arisanus* | | | 1,042 (154 Mb) | 978,588 nt | |
| *Macrocentrus cingulum* | | | 12,056 (128 Mb) | 65,089 nt | |
| *Microplitis demolitor* | | | 1,794 (241 Mb) | 1,139,389 nt | |
| *Venturia canescens* | | | 38,050 (234 Mb) | 117,151 nt | |

Supplementary Table 3. BUSCO analysis of parasitoid wasp genome assemblies

| **Species** | **Complete (% of total BUSCOs)** | **Duplicated (% of complete BUSCOs)** | **Fragmented (% of total BUSCOs)** | **Missing (% of total BUSCOs)** |
|---|---|---|---|---|
| *Adelognathus sp.* | 766 (46.2) | 3 (0.2) | 750 (45.2) | 142 (8.6) |
| *Aleiodes sp.* | 1428 (86.1) | 8 (0.5) | 191 (11.5) | 39 (2.4) |
| *Anoncus sp.* | 444 (26.8) | 3 (0.2) | 772 (46.6) | 442 (26.6) |
| *Dolichomitus sp.* | 473 (28.5) | 5 (0.3) | 778 (46.9) | 407 (24.6) |
| *Dusona sp.* | 1227 (73.6) | 6 (0.4) | 344 (20.7) | 87 (5.3) |
| *Eumacrocentrus americanus* | 1559 (93.2) | 13 (0.8) | 76 (4.6) | 23 (1.4) |
| *Lissonota sp.* | 863 (52.0) | 4 (0.2) | 651 (39.3) | 144 (8.7) |
| *Mesochorus sp.* | 1164 (70.2) | 10 (0.6) | 418 (25.2) | 76 (4.6) |
| *Meteorus sp.* | 1637 (98.7) | 7 (0.4) | 13 (0.8) | 8 (0.5) |
| *Odontocolon sp.* | 1170 (70.3) | 5 (0.3) | 366 (22.1) | 122 (7.3) |
| *Phanerotoma sp.* | 609 (36.7) | 2 (0.1) | 651 (39.3) | 398 (24.0) |

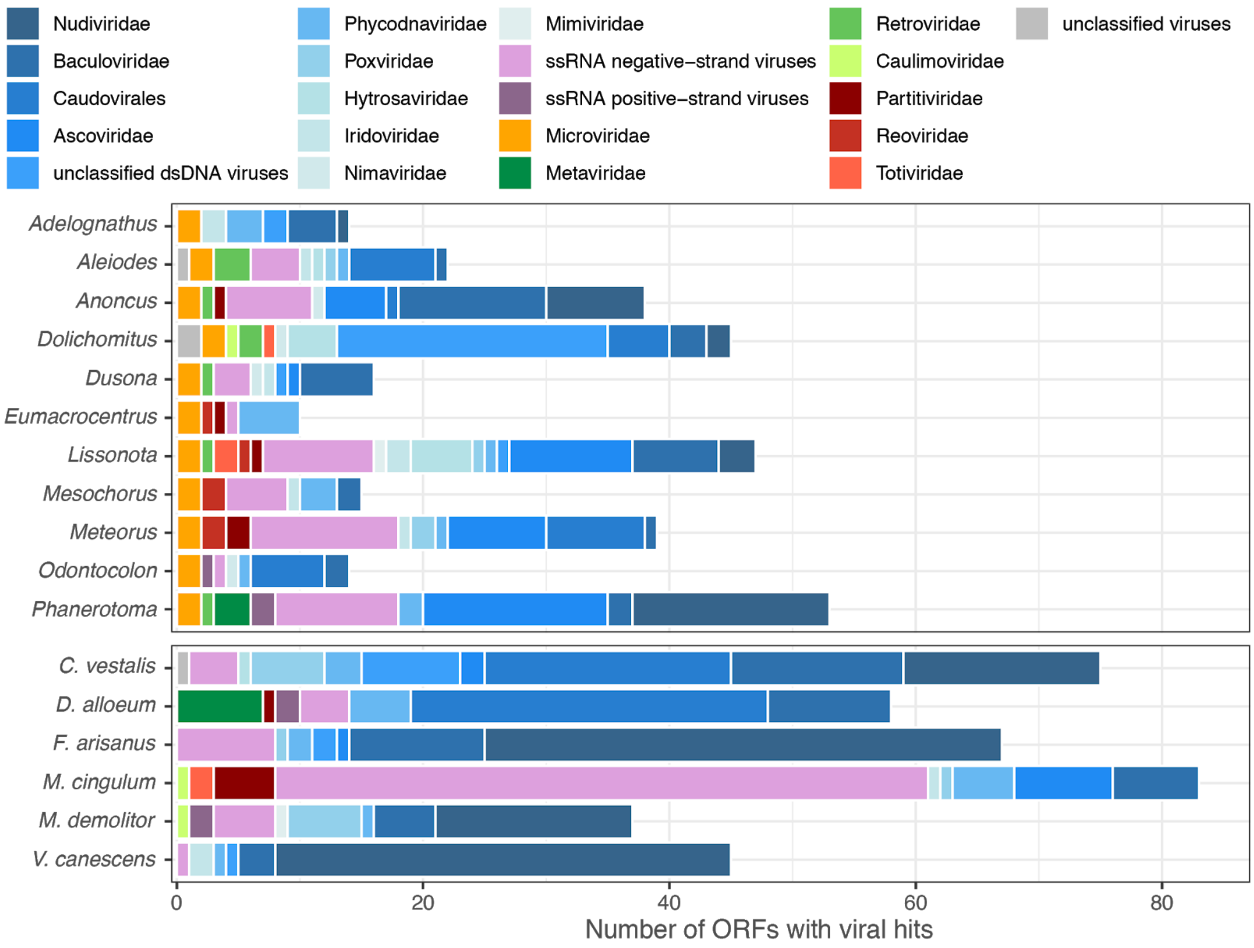


Supplementary Figure 1. Number of viral hits to different virus families for ORFs from wasp genome assemblies. New genome assemblies are shown in the upper panel, while previously published assemblies are shown below. Hits are colored by viral family and type: dsDNA viruses in blue, ssRNA viruses in purple, ssDNA viruses in orange, Ortervirales in greens, and dsRNA viruses in reds.

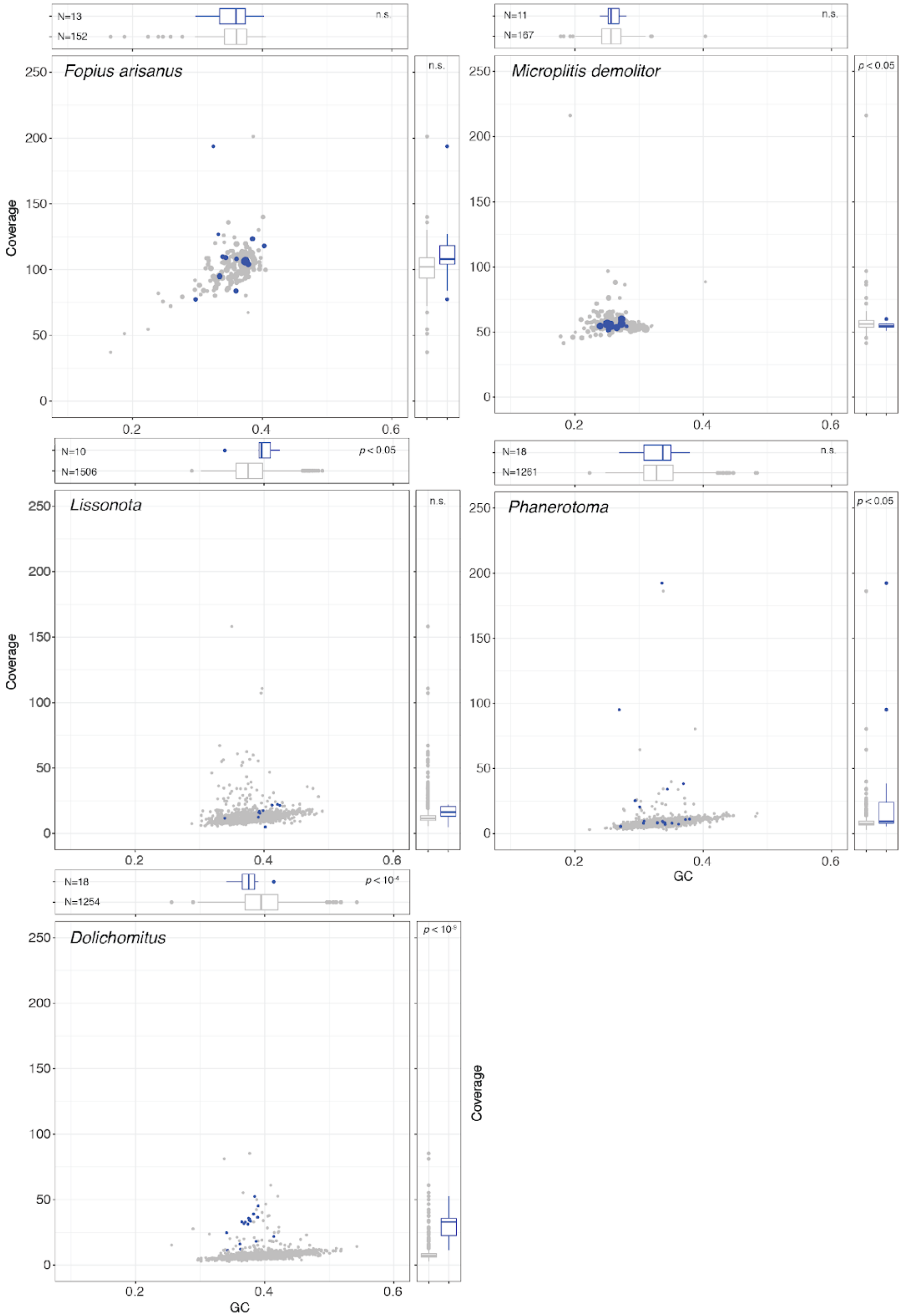


Supplementary Figure 2. Scatterplots and box plots of read coverage and GC content for contigs containing ORFs with viral hits (blue) compared to contigs containing BUSCOs (grey). In scatterplots, circles are sized based upon scaffold or contig length (ranging from 200bp to 2.7Mbp). Statistical tests (*t*-tests) were performed on $log_{10}$-transformed coverage values, and untransformed GC content values.
